# Phylogeography and Molecular Species Delimitation Reveal Cryptic and Incipient Speciation in Synchronous Flashing Fireflies (Coleoptera: Lampyridae) of Southeast Asia

**DOI:** 10.1101/632612

**Authors:** Wan F. A. Jusoh, Lesley Ballantyne, Chan Kin Onn

## Abstract

Synchronous flashing fireflies of the genus *Pteroptyx* are ubiquitous throughout Southeast Asia, yet, knowledge on its biodiversity and evolutionary history remains lacking. Recent studies have revealed notable population-level phylogeographic structure within the *P. tener* and *P. bearni* groups in Malaysia, suggesting that cryptic species may exist. Additionally, the close morphological and genetic affinity of the recently described species *P. balingiana* to *P. malaccae* has raised questions about its validity. In this study, we assembled the most densely sampled genetic dataset on *Pteroptyx* to-date to estimate a comprehensive phylogeny using mitochondrial and nuclear DNA and subsequently implemented a suite of distance-, phylogeny-, and coalescent-based species delimitation methods to characterize species boundaries within the *P. tener*, *P. bearni*, and *P. balingiana/P. malaccae* groups. Using a total evidence approach from multiple lines of evidence, we showed that populations of *P. tener* along the west coast of Peninsular Malaysia are sufficiently divergent from populations from the east coast and Borneo to warrant specific recognition, despite the absence of morphological differentiation. Conversely, divergence of *P. bearni* from Borneo and eastern Peninsular Malaysia, as well as *P. balingiana* from *P. malaccae* were modest and their distinction as separate species were ambiguous; consistent with incipient species in the gray zone of speciation. Overall, this study contributes to the limited but growing body of genetic work on Southeast Asian fireflies and underscores the urgent need to increase the breadth and depth of geographic, taxonomic, and genetic sampling to provide a deeper understanding of their biodiversity and evolutionary history.

## INTRODUCTION

Fireflies or lightning bugs are soft-bodied beetles of the family Lampyridae with the number of estimated species ranging from at least 2,000 to possibly 8,000 globally (Lloyd, 2008). Approximately 1,200 firefly species are known from tropical America (Faust, 2004), more than 400 species from Southeast Asia and the Indo-Pacific region (largely from the subfamily Luciolinae; Ballantyne, Lambkin, Boontop, & Jusoh, 2015; McDermott, 1966), while little information is available from tropical Africa (Lloyd, 2008).

Fireflies are best known for the ability of males to emit precise flashing patterns to attract females, although not all species are luminescent (Lloyd, 2002). The most spectacular of all flashing displays is the near-to-perfect synchronous flashing of fireflies from the genus *Pteroptyx* Olivier (subfamily Luciolinae), where species are known to occur in multitudes on trees and shrubs along tidal rivers of mangrove swamps (Jusoh, Ballantyne, Lambkin, Hashim, & Wahlberg, 2018). Its highly precise synchronous flashing and tree-swarming behaviour have attracted scientific as well public interests to explicate the mechanisms for their synchronicity (Buck, 2004), elucidate congregating and courting behaviour (Case, 2007; Lloyd, Wing, & Hongtrakul, 2006) and recently, develop as icon species for sustainable ecotourism and conservation (Khoo et al. 2012). However, although fireflies have been the focus of numerous ecological, physiological, and even biomedical research (Copeland & Moiseff, 2006; Ermentrout, 1991; Fraga, 2008; Gould Stephen J. & Subramani Suresh, 1988; Schena, Griss, & Johnsson, 2015), fundamental knowledge on its biodiversity and evolutionary history is still lacking and underscores the dire need for more intensive research.

According to recent studies (Ballantyne et al., 2015; Jusoh et al., 2018), *Pteroptyx* is an Oriental genus comprising 18 species distributed from Hong Kong, southwards throughout Southeast Asia including India (Madras) (Ballantyne, Fu, Shih, Cheng, & Yiu, 2011; Ballantyne & McLean, 1970) and the Philippines (Ballantyne, 2001). Most species in this genus are known to occupy mangroves, riparian, or coastal habitats (Ballantyne et al., 2011; Jusoh et al., 2018; Jusoh, Hashim, & Ibrahim, 2010). For instance, at least eight out of 12 species of *Pteroptyx* were recorded from mangroves throughout Malaysia (Jusoh et al., 2018), followed by four species in Thailand (Sartsanga, Swatdipong, & Sriboonlert, 2018), two to three species in Singapore (Chan, Ballantyne, & Goh, 2012), and one species in the Phillippines (Ballantyne et al., 2011; Jusoh et al., 2018). For centuries, species descriptions in the subfamily Luciolinae, which includes *Pteroptyx*, have been based purely on morphology. However, without morphological characters of the male genitalia and highly trained taxonomists, fireflies are difficult to identify to the species level (Ballantyne, 2012). Moreover, it is almost impossible to provide accurate identification of female and larval specimens, even to the genus level unless the specimens were collected in association with the male (Jusoh et al., 2018). Thus, additional methods and data streams are needed to facilitate species discovery in order to realize the true extent of firefly diversity in the region.

Recent developments in molecular sequencing technology have greatly improved our ability to detect, discover, and describe new species (Sites, Marshall, Jr, & Marshall, 2003; Wiens, 2007). Furthermore, comparing profiles of genetic divergences from informative markers have been shown to be an effective way to identify species and screen for taxonomic incompatibilities and potential new species (Chan & Grismer, 2019; Chan, Grismer, & Brown, 2018; Fouquet et al., 2007; Hebert, Cywinska, Ball, & DeWaard, 2003; Padial & De La Riva, 2007; Vences, Thomas, Bonett, & Vieites, 2005; Vences, Thomas, Meijden, Chiari, & Vieites, 2005; Vieites et al., 2009). Among the genetic markers implemented for these purposes, the DNA barcode based on the *CO1* mitochondrial gene is one of the most widely used, especially among invertebrates. Although it was initially developed for species identification (Hebert et al., 2003), this robust marker has also been used effectively for species discovery and delimitation (Cao et al., 2016; Hebert & Gregory, 2005; Hubert & Hanner, 2016; Sheth & Thaker, 2017; Yang & Rannala, 2017).

DNA sequence data for fireflies, in particular the *CO1* gene, have recently been accumulating but a well-characterized profile of genetic variation and a justified threshold for species boundaries have yet to be determined. The first study that implemented DNA barcoding on several species of *Pteroptyx* in Malaysia found congruence between morphology and molecular data (Jusoh, Hashim, Sääksjärvi, Adam, & Wahlberg, 2014). However, molecular results also revealed considerable phylogeographic structure (Fig. 2 in Jusoh *et al.*, 2014), suggesting that the genus *Pteroptyx* could potentially harbour cryptic or undiscovered species. In particular, *Pteroptyx tener* Olivier was clustered into three geographically isolated clades: along the east and west coast of Peninsular Malaysia (PM east and PM west), and Borneo (Fig. 1). In Thailand, a population of *P. tener* was discovered for the first time in 2015, which prompted the hypothesis that geographic polymorphisms may exist (Sriboonlert, Swatdipong, Wonnapinij, E-Kobon, & Thancharoen, 2015). Subsequent examination of morphological characters did not reveal differences that were congruent with the geographically circumscribed haplotypes (Jusoh et al., 2018). *Pteroptyx bearni* Olivier on the other hand, exhibited subtle differences in colouration and was geographically and genetically clustered into two clades: PM east and Borneo (Jusoh et al., 2014).

**Fig. 1.**
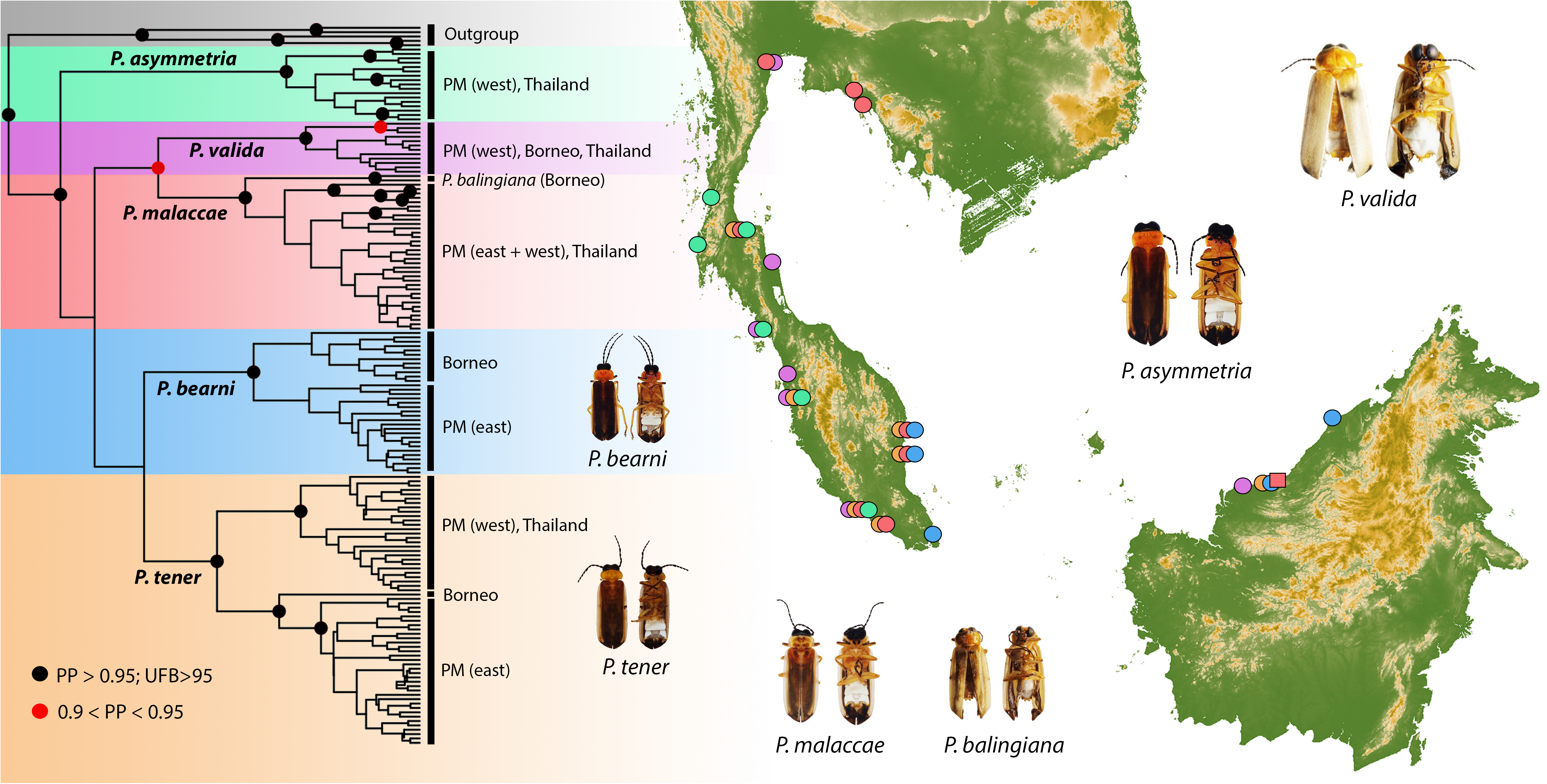
Geographic distribution of *Pteroptyx* samples used in this study and the BEAST phylogeny inferred from a concatenated sequence matrix consisting of 2,119 bps of the *CO1* mitochondrial and CAD nuclear genes. Circles at nodes denote Bayesian posterior probabilities and ML Ultrafast Bootstrap support values. Nodes without circles have low support (PP<0.9; UFB<95). Species-level clades are color-coded to match locality points on the distribution map. Inset figures show dorsal and ventral views of male representative species from the ingroup.

*Pteroptyx malaccae* (Gorham, 1880) is a widely distributed species in mangroves throughout Southeast Asia (Ballantyne, 1987; Ballantyne & McLean, 1970). In Malaysia, *P. malaccae* occurs in small populations, usually in sympatry with *P. tener*, which appears in much bigger congregations (Foo & Dawood, 2017; Jusoh et al., 2018). Meanwhile in Thailand, *P. malaccae* forms mass congregations in trees along the riverbanks in sympatry with *Pteroptyx valida* Olivier (Prasertkul, 2018; Sartsanga et al., 2018). Jusoh *et al.* (2018) hypothesized that *P. malaccae* is a morphologically variable species complex that requires further investigation after four morphologically distinct groups (Ballantyne, 2001) were detected using a combination of morphological and genetic data. The same analysis, although less resolved, also revealed that *P. malaccae* formed a sister relationship with a newly described species, *Pteroptyx balingiana* Jusoh. The authors previously argued that the latter species (although misidentified as *Pteroptyx gelasina* Ballantyne, see Jusoh et al. 2018) could be “isolated from the *P. malaccae* by distinct geographically structured variation in the DNA barcodes” (Jusoh et al., 2014).

Both previous studies (Jusoh et al., 2018, 2014) suggested that the spatio-genetic structure of populations within *Pteroptyx tener, P. bearni*, and *P. malaccae/P. balingiana* could represent complexes containing cryptic and undescribed species, but no explicit species delimitation analyses have been performed to validate those hypotheses. In this study, we assembled the most densely sampled genetic dataset of *Pteroptyx* to-date to estimate a comprehensive phylogeny using mitochondrial and nuclear DNA. We then employed a suite of distance-, phylogeny-, and coalescent-based species delimitation methods to test whether 1) geographically structured populations in *P. tener* and *P. bearni* represent cryptic species; and 2) *Pteroptyx balingiana* is sufficiently genetically distinct from *P. malaccae* to warrant specific recognition.

## METHODS

### Taxon coverage and geographic sampling

We used sequence data from published studies that are publicly available on GenBank and constructed a dataset consisting of 157 sequences of the mitochondrial *Cytochrome c oxidase subunit 1* (*CO1*) gene and 39 sequences of the nuclear protein coding gene, *carbamoylphosphate synthetase* (CAD; Supplementary information, Table S1). Apart from the focal taxa (*Pteroptyx malaccae, P. balingiana, P. tener*, and *P. bearni*), we also included other closely related taxa such as *Pteroptyx asymmetria* Ballantyne and *P. valida*, while outgroup samples include *Pteroptyx galbina* Jusoh, *Pteroptyx testacea* (Motschulsky), *Colophotia praeusta* (Eschscholtz) and *Colophotia brevis* Olivier. The focal taxa were sampled from 23 unique localities across Thailand and Malaysia (Fig. 1). The *CO1* gene comprises three separate fragments, which include the prominent “Folmer region” of a 665 bp fragment located at the 5’ end of the *CO1* gene (Folmer, Black, Hoeh, Lutz, & Vrijenhoek, 1994), a ~540 bp fragment at the 3’ end (Villalba, Lobo, Martin-Piera, & Zardoya, 2002), and a long fragment (~1,310 bp) that overlaps with the first two fragments (Sartsanga et al., 2018). Samples that contain more than one of these fragments were combined to form a single contig. Sequences from the second *CO1* fragment which were amplified using the primers *CO1-Sca (F)/CO1-Sca* (*R*) (Villalba et al., 2002) overlapped with the *Cytochrome c oxidase subunit 2* (*CO2*) gene as well as a *tRNA-Leu* and was removed prior to sequence alignment. Both *CO1* and *CAD* sequences were checked for stop codons and aligned separately using MUSCLE with default parameters performed on MEGA-X version 10.0.5 (Kumar, Stecher, Li, Knyaz, & Tamura, 2018). Alignment files are available as Supporting information.

### Phylogenetic analyses

We used the program IQ-TREE v1.6 (Nguyen, Schmidt, Von Haeseler, & Minh, 2015) to estimate a maximum likelihood (ML) phylogeny, while a Bayesian phylogeny was inferred using BEAST v2.5 (Bouckaert et al., 2014). For the ML phylogeny, sequences were partitioned by gene and the best-fit model of DNA evolution for each partition was determined using ModelFinder (Kalyaanamoorthy, Minh, Wong, Von Haeseler, & Jermiin, 2017). Branch support was assessed with 5,000 bootstrap replicates using Ultrafast Bootstrap Approximation (UFB; Hoang et al. 2017). UFB values above 95 were considered well supported. The BEAST analysis was implemented through the CIPRES portal (Miller, Pfeiffer, & Schwartz, 2010) and the best-fit substitution model for each partition was estimated via model averaging using the bModelTest plugin in BEAST (Bouckaert & Drummond, 2017). A relaxed log-normal and Yule model was used as the molecular clock and tree priors respectively, while all other priors were set to default values. We executed two separate MCMC chains at 50 million generations each and checked for convergence using the program Tracer v1.6 (Rambaut, Drummond, Xie, Baele, & Suchard, 2018). Sampled trees from both MCMC runs were combined using *logcombiner* and a maximum clade credibility tree was constructed using *treeannotator* with a burn-in of 10% and nodes summarized using mean heights.

### Species delimitation

*Genetic distance and ABGD*.—we calculated uncorrected p-distances within and between taxa using MEGA-X (Kumar et al., 2018) to obtain a profile of intra- and interspecies divergence thresholds. Subsequently, the Automatic Barcoding Gap Discovery (ABGD) method was used to detect breaks between intraspecific and interspecific diversity known as the barcode gap (Puillandre, Lambert, Brouillet, & Achaz, 2012). The ABGD uses pairwise distances to determine divergence between sequences and does not require a priori specification of divergence thresholds. Because ABGD analyzes single-locus data and has been shown to be effective with the CO1 gene, we performed this analysis on the CO1 alignment. The analysis was performed through the web-server (http://wwwabi.snv.jussieu.fr/public/abgd/abgdweb.html) using default settings and the Kimura 2-parameter (K80) distance model (Ratnasingham & Hebert, 2013).

*bGMYC*.—compared to ABGD, the Bayesian implementation of the general mixed Yule-coalescent (bGMYC) is a non-distance-based method that does not rely on similarity threshold parameters. It models the Yule and coalescent processes on an ultrametric tree to determine the transition between intra and interspecific divergences. To account for potential errors in phylogenetic estimation and uncertainty in model parameters, this method integrates over uncertainty in tree topology and branch lengths via MCMC (Reid & Carstens, 2012). As input, we used 100 randomly selected trees that were sampled from the posterior distribution of the BEAST analysis. For each tree, we ran the MCMC sampler for 50,000 generations with a burnin of 40,000, retaining 10,000 post burn-in generations with a thinning interval of 100. We then assessed a range of probability thresholds ranging from conservative (0.05) to moderate (0.25) and liberal (0.5) delimitation schemes. The analysis outputs results in the form of posterior probabilities that sequences are conspecific. Hence, we consider populations with PP<0.05 as strong support, 0.1< PP< 0.05 as moderate support, and PP > 0.1 as weak support for splitting.

*mPTP*.—the multirate Poisson tree process (mPTP) is also a phylogeny-aware method that also does not rely on similarity threshold parameters. Similar to bGMYC, it models the transition between intra- and interspecific divergence that is determined by the coalescent and speciation parameters, where intraspecific branching events are expected to be more frequent compared to among species (Kapli et al., 2017; Zhang, Kapli, Pavlidis, & Stamatakis, 2013). However, mPTP differs from bGMYC in modelling speciation and coalescent events relative to numbers of substitutions instead of time (Tang, Humphreys, Fontaneto, & Barraclough, 2014). It takes into account the evolutionary relationships of sequences and uses the number of accumulated expected substitutions between subsequent speciation events to model the branching process. We used the ML phylogeny as the input tree and confidence of delimitation schemes were assessed using two independent MCMC chains at 5,000,000 generations each. Support values represent the fraction of sampled delimitations in which a node was part of the speciation process.

*BPP*.—the BPP analysis uses a Bayesian modelling approach to estimate posterior probabilities of species assignments using gene trees, while taking into account uncertainties in the coalescent process (Yang & Rannala, 2010). The A10 analysis (species delimitation using a fixed guide tree) was implemented using relationships derived from phylogenetic analyses as a guide tree. We used a diffuse prior of α =3 for both θ and τ priors and the corresponding β parameter was adjusted according to the mean (m) estimate of nucleotide diversity (for θ) and node height (for τ) using the equation m=β/(α−1), for α>2 (Flouri, Jiao, Rannala, & Yang, 2018). The mean root node height was obtained from the BEAST phylogeny. The use of empirically-estimated priors has been shown to provide more reliable results compared to priors selected using non-objective methods (Chan & Grismer, 2019). To accommodate uncertainty in the guide tree, we also performed the A11 analysis (joint species delimitation and species-tree estimation). The MCMC was set to 100,000 samples with burnin=10,000 and sample frequency=5. Convergence was assessed by comparing the consistency of posterior distributions (Flouri et al., 2018; Leaché, Zhu, Rannala, & Yang, 2018; Ziheng, 2015). Posterior probabilities, PP ≥ 0.95 were considered highly supported, 0.90 ≤ PP < 0.95 were considered moderately supported, whereas PP < 0.9 were considered weakly supported.

*Heuristic gdi*.—because Bayesian model selection in BPP can sometimes oversplit species by detecting population splits as species divergence (Leaché et al., 2018; Sukumaran & Knowles, 2017), we used the heuristic genealogical index index (*gdi*) that has been shown to produce more accurate results (Chan & Grismer, 2019; Jackson, Carstens, Morales, & O’Meara, 2017a; Leaché et al., 2018). First, the A00 analysis in BPP was implemented to generate posterior distributions for the parameters τ and θ using the same empirically-derived priors. Four separate runs were performed to ensure convergence and converged runs were combined to generate posterior distributions for the MSC parameters that were subsequently used to calculate the *gdi* following the equation: *gdi* = 1 – e^−2τ/θ^ (Jackson, Carstens, Morales, & O’Meara, 2017b; Leaché et al., 2018). Population A is distinguished from population B by using 2τ_AB_/θ_A_, while 2τ_AB_/θ_B_ is used to differentiate population B from population A. Populations are considered distinct species when *gdi* values are >0.7, while low *gdi* values <0.2 indicate that populations belong to the same species. Values of 0.2> *gdi* < 0.7 indicate ambiguous species status (Jackson et al., 2017b; Pinho & Hey, 2010).

## RESULTS

### Phylogenetic analyses

Both ML and Bayesian phylogenies produced congruent topologies with high support across most major nodes (Figs. 1, S1, S2). Known species also formed monophyletic clades with high support. *Pteroptyx valida, P. asymmetria*, and *P. malaccae* did not show geographically circumscribed genetic sub-structuring, with rampant mixing occurring between individuals from Peninsular Malaysia (PM) and Thailand. *Pteroptyx balingiana* was reciprocally monophyletic with *P. malaccae* with high support, whereas the *P. bearni* and *P. tener* clades contained multiple populations that were distinctly structured according to geography. *Pteroptyx bearni* formed two distinct clades (Borneo and PM), while *P. tener* was structured into three relatively divergent clades. The Bornean population of *P. tener* formed a sister relationship with populations from eastern PM and this clade (Borneo + PM east) was in turn reciprocally monophyletic with populations from western PM (Fig. 1). Using this phylogeographic framework, we defined the following geographically circumscribed and reciprocally monophyletic population pairs as candidate species for downstream species delimitation analyses: i) balingiana vs. malaccae; ii) bearni Borneo vs. bearni PM; iii) tener Borneo vs. tener PM east; and iv) tener PM west vs. tener Borneo+PM east (Fig. 1).

### Genetic divergence and species delimitation

*Genetic distance*.—the interspecies genetic divergence between the sister lineages *P. bearni/ P. tener* and *P. valida/P. malaccae* were high (6.3–8.9% and 8–10.5% respectively), while divergences among populations of the same species were low (<3%). For the candidate species, divergences were slightly higher compared to intrapopulation distributions, except for tener PM west vs. tener Borneo + tener PM east which was higher (3.15–5.49%), but still well below interspecific divergences (Fig. 2).

**Fig. 2.**
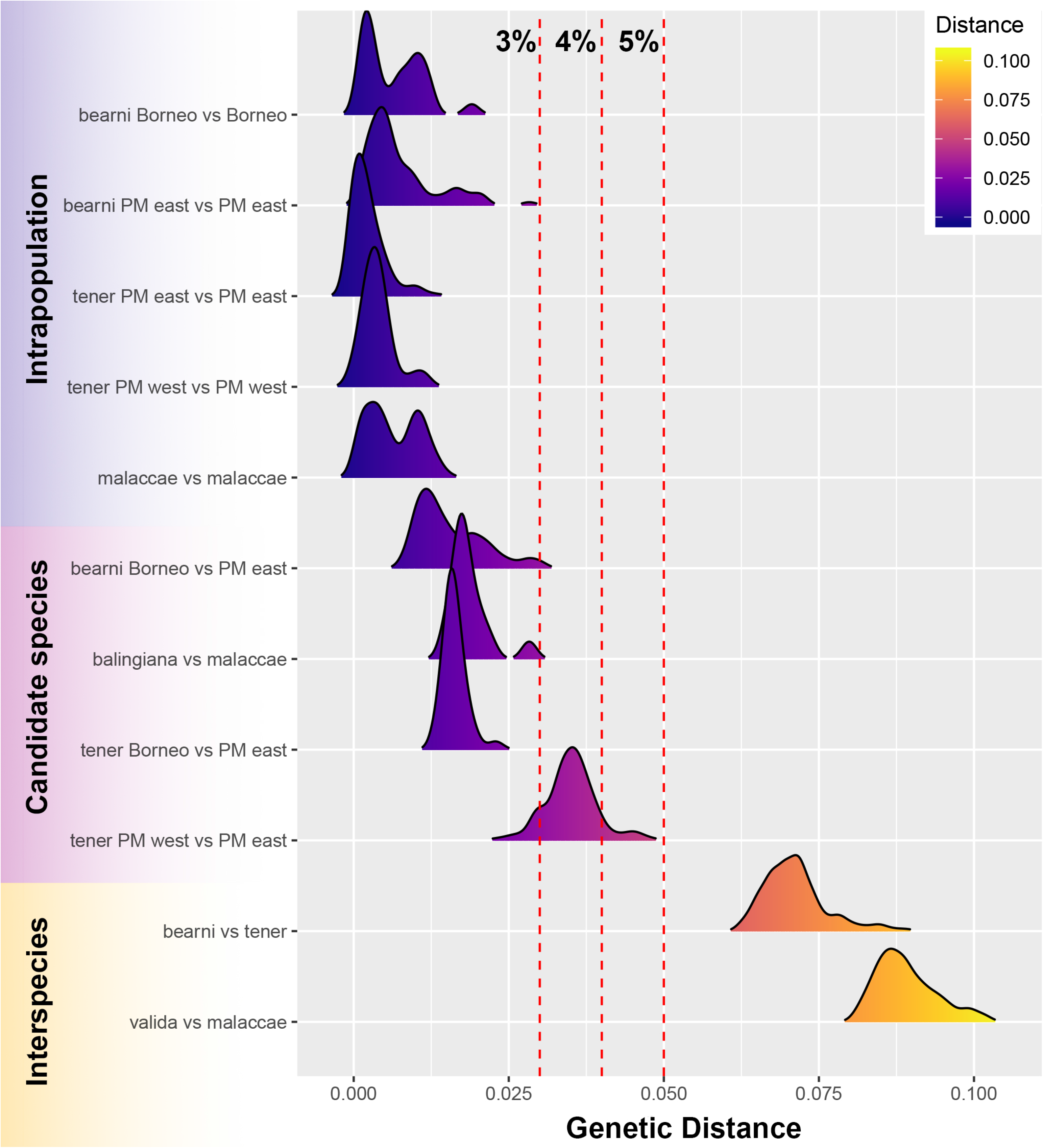
Distribution of intra and interspecific genetic distances based on the *CO1* mitochondrial gene. Candidate species represent reciprocally monophyletic and geographically circumscribed populations that were subjected to downstream species delimitation analyses. Dotted red lines represent the 3%, 4%, and 5% genetic distance thresholds.

*ABGD*.—a total of seven partitions were inferred with prior maximal intraspecific distances (P) ranging from 0.001 to 0.02 (Fig. 3; Table 1). For the initial run, the number of delimited species rapidly plateaued at 10 species. This delimitation scheme lumped balingiana with malaccae, bearni Borneo with bearni PM, and tener Borneo with tener PM east. However, tener PM west was delimited as a distinct species from tener Borneo + tener PM east (Table 2). At P=0.0077, the recursive run also recovered the same 10 species but at P=0.0046 (partition 4), tener Borneo was further split from tener PM east. Partitions 1–3 of the recursive run produced 13–26 species, which we consider as erroneous as some of the delimited groups were not monophyletic.

**Fig. 3.**
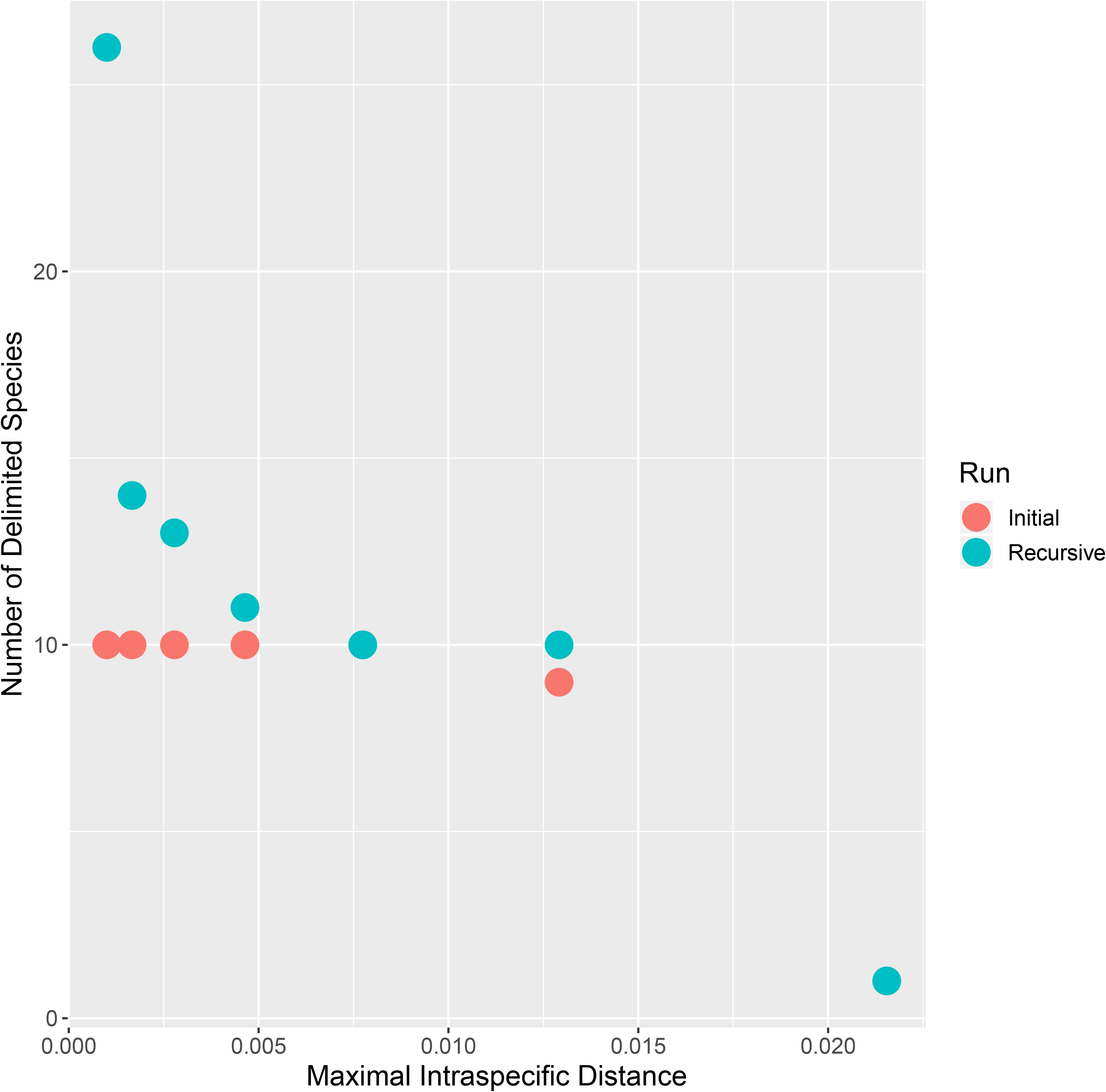
Results of the initial and recursive runs of ABGD and the number of delimited species at the associated maximal intraspecific distance (P).

**Table 1.** Partitions, number of species (initial run followed by recursive run in parenthesis), and corresponding prior maximal distance from the ABGD analysis using the Kimura (K80) distance model and a relative gap with of X=1.5.

|  | Number of species | Prior max. distance |
| --- | --- | --- |
| Partition 1 | 10(26) | 0.001 |
| Partition 2 | 10(14) | 0.001668 |
| Partition 3 | 10(13) | 0.002783 |
| Partition 4 | 10(11) | 0.004642 |
| Partition 5 | 10(10) | 0.007743 |
| Partition 6 | 9(10) | 0.012915 |
| Partition 7 | 1(1) | 0.021544 |

**Table 2.** Summary of the species delimitation results on the candidate species.

|  | balingiana vs.<br>malaccaae | bearni Borneo<br>vs. bearni PM | tener Borneo<br>vs. tener PM east | tener Borneo+PM east<br>vs. tener PM west |
| --- | --- | --- | --- | --- |
| P- distance (%) | 1.48–2.92 | 0.84–5.00 | 1.39–2.29 | 3.15–5.49 |
| ABGD □ | Lump/split | Lump/split | Lump/split | Split |
| mPTP □ | Low (0.16) | Low (0.13) | Low (0.67) | High (1.0) |
| bGMYC □ | Low (0.29) | Low (0.23) | Low (0.23) | Moderately High (0.08) |
| BPP (A10) □ | High (1.0) | High (1.0) | High (1.0) | High (1.0) |
| BPP (A11) □ | High (1.0) | High (1.0) | High (1.0) | High (1.0) |
| gdi □ | High | Uncertain | Uncertain | High |

□ Combined results from partitions 4–6 that delimited 9–10 species. See Results for more details.
□ Support values denote posterior probabilities that lineages are separate species.
$PP > 0.95$ =High; $0.9 < PP < 0.95$ = Moderate; $PP < 0.9$ = Low.
□ Support values denote posterior probabilities that lineages are conspecific. Hence,
lineages are highly supported as separate species if $PP < 0.05$ , while $0.1 > PP > 0.05$
= moderate support; $PP > 1.0$ = Low support.
□ Average $gdi > 0.7$ highly supports lineages as separate species; $0.2 < gdi < 0.7$
denotes uncertain species status, while $gdi < 0.2$ denotes lineages are the same
species.

*bGMYC*.—the distribution of the coalescence to Yule branching rate ratios were well above zero, indicating that the model is a good fit to the data. The results showed low support for the splitting of balingiana/malaccae (conspecificity PP=0.29), bearni Borneo/PM (PP=0.23), and tener Borneo/PM east (PP=0.5). On the other hand, the splitting of tener Borneo + tener PM east/tener PM west was moderately supported (PP=0.08; Table 2). At the most conservative threshold of 0.05, the analysis lumped balingiana with malaccae, and all populations of *P. bearni* and *P. tener*. At 0.25, balingiana remained lumped with malaccae, while tener Borneo was still lumped with tener PM east. However, bearni Borneo was split from bearni PM and tener PM west was split from tener Borneo + tener PM east. The most liberal threshold of 0.5 further split balingiana from malaccae and tener Borneo from tener PM east. Certain samples of *P. malaccae* from Thailand were also split from other *P. malaccae* (including other Thai samples) at this threshold, indicating that this threshold was too liberal at splitting species.

*mPTP*.—the mPTP analysis did not support the splitting of balingiana and malaccae (PP=0.16); bearni Borneo and bearni PM (PP=0.13); or tener Borneo and tener PM east (PP=0.67); but highly supported the split between tener PM west and tener Borneo + tener PM east (PP=1.0).

*BPP* and *gdi*.—all candidate populations were delimited as distinct species with high support (posterior probability=1.0) in the BPP A10 and A11 analyses (Table 2). The heuristic *gdi* analysis provided moderately high but uncertain support for the split between balingiana and malaccae; tener Borneo and tener PM east; and bearni Borneo and bearni PM. The split between tener PM west and tener Borneo + tener PM east was highly supported (Fig. 4; Table 2).

**Fig. 4.**
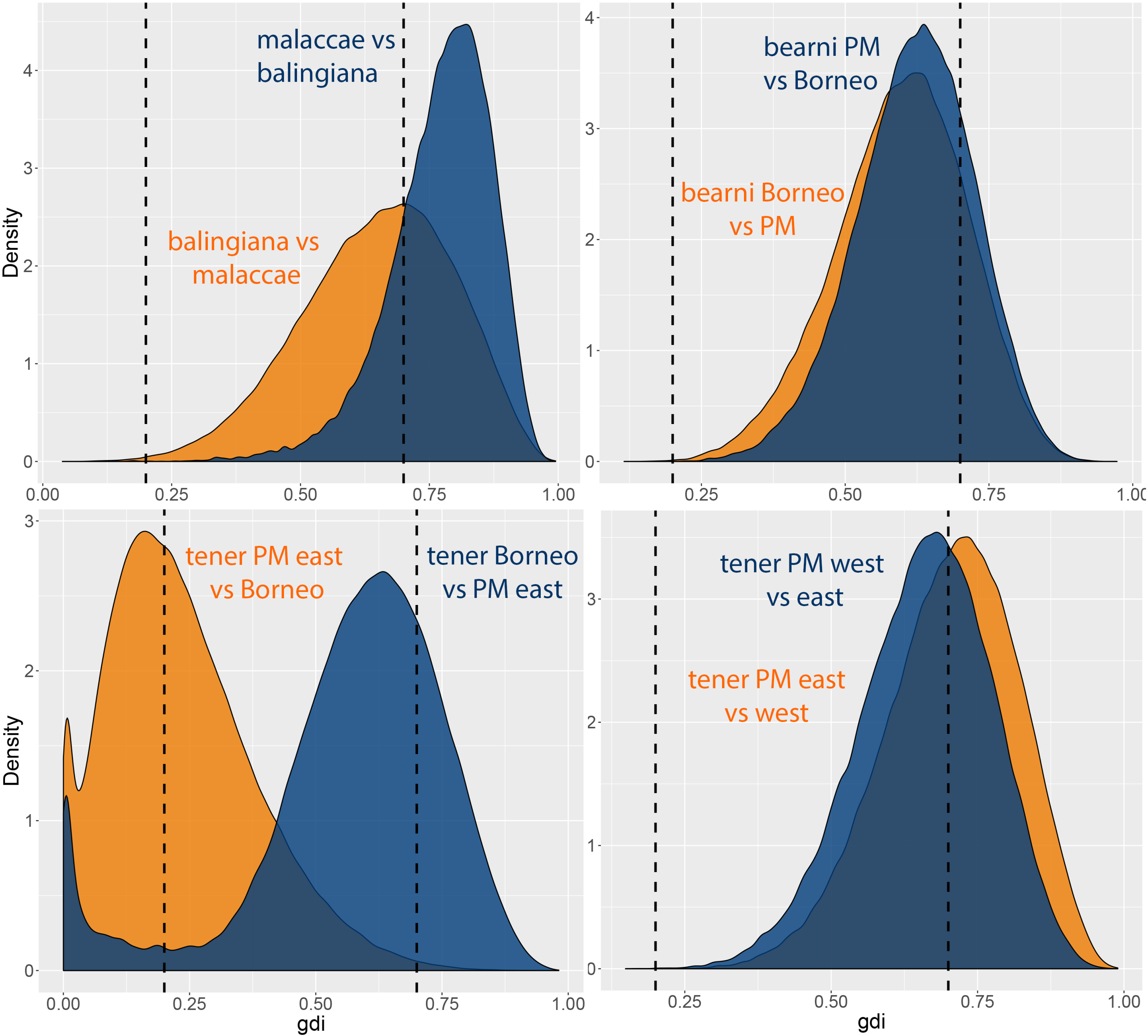
Density plots of *gdi* values with vertical dotted lines representing the 0.2 and 0.7 thresholds. Populations are considered distinct species if *gdi* >0.7, while *gdi* <0.2 indicate that populations belong to the same species. Values of 0.2> *gdi* < 0.7 indicate ambiguous species status.

## DISCUSSION

### Phylogenetic relationships

Phylogenetic relationships in Luciolinae were exclusively based on ~400 morphological characters (Ballantyne & Lambkin, 2013; Ballantyne et al., 2015) until Jusoh et al. (2018) made the first attempt to combine and jointly analyze the morphological data with a small subset of molecular data from 25 taxa (total = 158 taxa) using Bayesian Inference (BI) and Maximum Parsimony (MP). Although taxon sampling was limited, the molecular data recovered a similar topology with the larger morphological matrix, but the combined morphological and molecular dataset analysis produced different relationships. Within the *Pteroptyx* clade, the MP analysis (Figure 2, part 2 in Jusoh et al. 2018) revealed that *P. balingiana* formed a clade with *Pteroptyx macdermotti* McLean and *P. gelasina* and this clade was reciprocally monophyletic with *P. malaccae* (the BI analysis was unresolved). This does not conflict with the topology from this study as no molecular samples of *P. macdermotti* and *P. gelasina* were available for analysis. However, the phylogenetic position of *P. valida* and *P. asymmetria* were not concordant with results from our analyses. Our analysis recovered *P. asymmetria* as the basal lineage with regard to *P. valida, P. malaccae, P. bearni*, and *P. tener*, but Jusoh et al (2018) recovered it within the *P. bearni* + *P. tener* subclade, which was in turn sister to the *P. valida* clade. Based on morphology, *P. asymmetria* is more affine to *P. tener* and *P. bearni*. Our study recovered *P. valida* as the sister lineage to the *P. malaccae* + *P. balingiana* subclade albeit with moderate support. Unfortunately at this point, we are still unable to resolve the species relationships of Malaysia *Pteroptyx* with any amount of certainty and more data in the form of genetic loci and taxon representation will be needed to provide more robust inferences.

### Species boundaries and systematics

Using a suite of species delimitation analyses that are based on different models and assumptions, we showed that *Pteroptyx* from Southeast Asia runs the gamut of the speciation continuum and exhibits strong phylogeographic patterns that are in some cases incongruent with morphological differentiation. Populations of *P. tener* from Borneo + PM east showed high genetic divergence and was highly supported as distinct from the PM west population across all analyses. Furthermore, sampling for each of these populations were robust and represented by multiple individuals. As such, the results can be considered robust and the specific distinction between these lineages can be posited with confidence. However, there are no reliable diagnostic characters that distinguish these lineages (see Fig. 5A; Jusoh et al., 2018), indicating that they constitute cryptic species (Bickford et al., 2007).

**Fig. 5.**
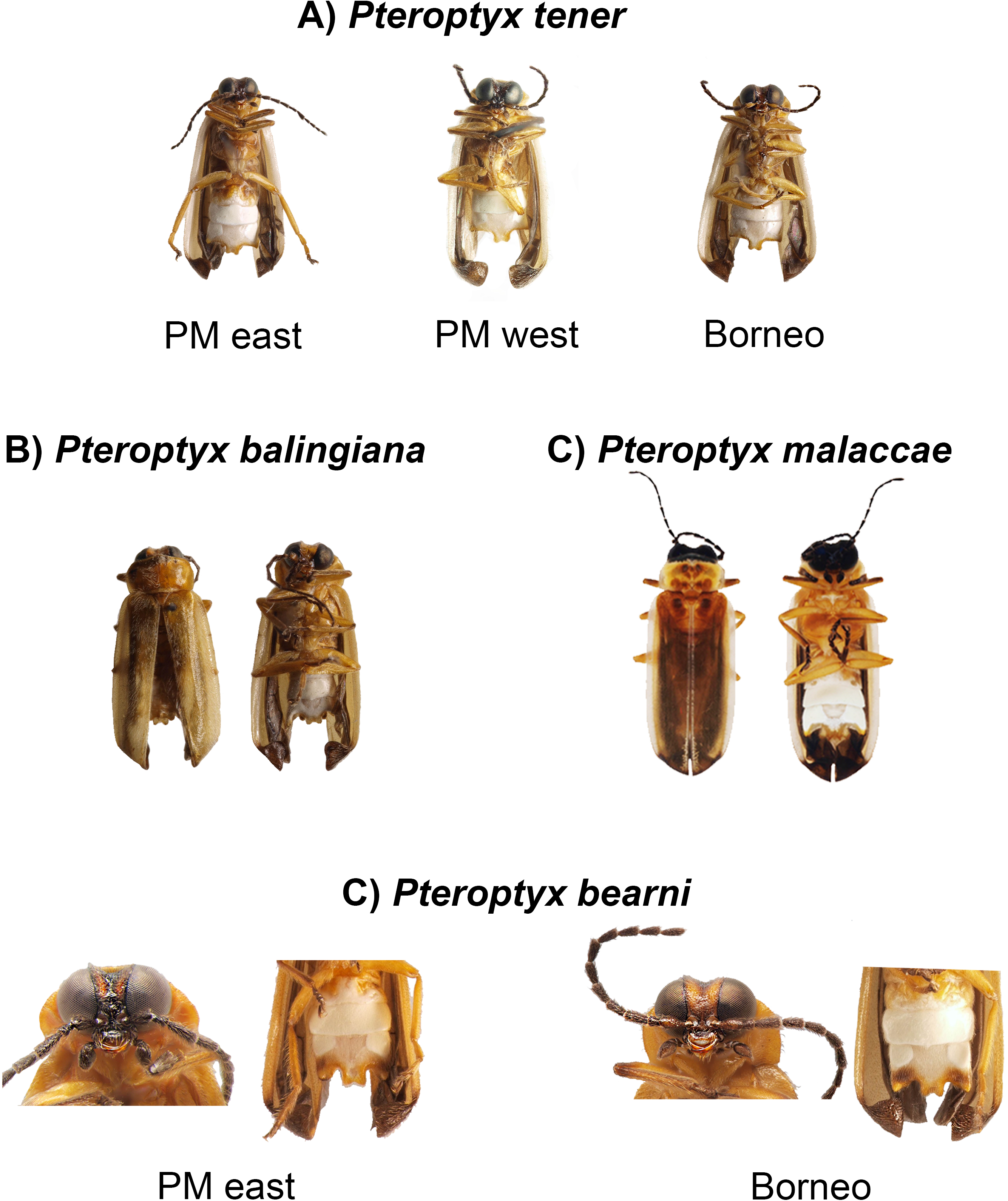
**A.** Ventral view of *Pteroptyx tener* from PM east, PM west, and Borneo; **B and C.** dorsal and ventral view of *P. balingiana* and *P. malaccae* (note the difference in the shape of the light organ); **D.** *P. bearni* from PM east and Borneo can be differentiated by the colouration of the head but both populations have similar light organ structures.

*Pteroptyx balingiana* was described as a separate species from *P. malaccae* on the basis of morphological and molecular differentiation (Jusoh et al., 2018). Morphologically, *P. balingiana* can be distinguished from *P. malaccae* by structural differences in the light emitting organ (Fig. 5B,C). More in-depth analyses performed here demonstrated that genetic divergence between the two taxa was moderate and their status as distinct species was not unequivocally supported by species delimitation analyses. Nevertheless, the data also did not strongly support the lumping of these taxa. In light of the following lines of evidence: i) presence of morphological differences; ii) allopatry; iii) genetic differentiation (albeit modest); these taxa clearly represent independently evolving metapopulation lineages that can be considered as distinct species under the General Lineage Concept of Species (de Queiroz, 2007). As such, we retain the current taxonomic status of *P. balingiana* and *P. malaccae* as distinct species until additional data suggest otherwise.

Similarly, species delimitation results for populations of *P. bearni* from Borneo/ PM and populations of *P. tener* from Borneo/PM east were also largely uncertain. According to the morphological examination of Jusoh et al. (2018) which consist of freshly preserved specimens, individuals of *P. bearni* from PM and Borneo can be distinguished by the colouration on the pronotum and head (Fig. 5D). However, these differences were not observable in preserved specimens. No morphological differences were detected between populations of *P. tener* from Borneo and eastern PM (Fig. 5A), but this could be due to the limited number of Bornean samples. Based on the relatively shallow bipartitions of these lineages and the absence or subtle differences in morphology, we interpret them as being in the early stages of speciation where insufficient time has elapsed for considerable genetic differences to accumulate. Demonstrably, the distribution of genetic divergences show slight but clear shifts away from population-level variation, but have yet to attain the levels of divergence observed between species (Fig. 2). We therefore consider these lineages as incipient species that have begun to diverge, but remain in the gray zone of the speciation continuum (Feder, Egan, & Nosil, 2012; Nosil & Feder, 2012; Roux et al., 2016). Because evidence was not compelling, we refrain from splitting these populations into separate species pending further investigation.

### Caveats, limitations, and future directions

Both mPTP and bGMYC methods seek to identify the transition point between the speciation and coalescent process, hence adequate taxon representation is needed to provide robust estimates. Consequently, populations represented by singletons or doubletons can affect the accuracy of the analysis (Kapli et al., 2017; Reid & Carstens, 2012). While BPP has been shown to be less sensitive to taxon rarity or low numbers of loci (Yang & Rannala, 2017), this analysis is also known to oversplit species due to its inability to differentiate between population structure and species divergence under a protracted speciation model (Chan & Grismer, 2019; Leaché et al., 2018; Sukumaran & Knowles, 2017). This could explain the unrealistically high support for splitting every one of the focal populations, which is in conflict with results from the other species delimitation analyses. Therefore, the BPP analyses and results pertaining to populations with singletons or doubletons should be treated with caution.

The limited number of genes that was available for analysis could also potentially affect the results. Unfortunately, there is a severe lack of nuclear gene representation in public databases. Furthermore, additional studies are required to identify informative genes that can be useful for phylogenetics and species delimitation in fireflies, as this has yet to be determined. There is also a dearth of taxonomic representation in molecular samples where only eight out of 18 species of *Pteroptyx* have ever been sequenced, rendering the phylogenetic relationships of this genus uncertain. Morphologically, *Pteroptyx* is a genus with three subdivisions: Group I (deflexed elytra + meta femoral comb (MFC) + bipartite light organ (BLO)) (see Fig. 5); Group II (non-deflexed elytra + MFC + entire light organ (ELO)); and Group III (non-deflexed elytra + no MFC + BLO; see Jusoh et al. 2018). This study only dealt with taxa from Group I or commonly known as the bent-winged fireflies, with poor representative from the other groups (Groups II = *P. galbina*, III = *P. testacea*). Future phylogenetic studies should ideally include key taxa that are closely related to *Pteroptyx* such as *Medeopteryx* Ballantyne and *Pyrophanes* Olivier. Another species that is morphologically similar to *P. malaccae* and *P. balingiana* is *P. gelasina*, which is only known from Sabah, Borneo. Unfortunately, no molecular data is available for this species. Therefore, apart from increasing the genetic sampling of *P. balingiana*, acquiring molecular data for *P. gelasina* is also crucial to determine the species boundaries and phylogeographic structure of this species complex. While missing taxa could affect the topology of the overall phylogeny, we do not anticipate that it will significantly disrupt our species delimitation results as those were performed at the population level (i.e. the populations within the *P. malaccae, P. bearni*, and *P. tener* groups), which are not affected by uncertainty in relationships at the species level.

### Biogeography

Due to the limited number of genes and the absence of calibration points, we were unable to anchor the phylogeny onto a temporal framework of absolute time and this prevented us from providing an estimate on diversification times. However, based on the relatively shallow divergences among populations from PM and Borneo, we hypothesize that cyclical glaciations during the Pleistocene (Hall, 2013) could have facilitated the vicariance of these populations, resulting in the fragmented present-day distribution on their respective land masses (Erik, 2003; Lim et al., 2017; Slik et al., 2011; Wurster et al., 2010). Within Peninsular Malaysia, the east-west pattern of lineage separation has been demonstrated in numerous co-distributed taxa (Chan, Abraham, Grismer, & Grismer, 2018; Chan et al., 2017, 2014; Grismer et al., 2013, 2015). The isolation of those taxa were hypothesized to be caused by the relatively central and contiguous Titiwangsa mountain range, which acts as an insurmountable physical barrier that separates habitats along the eastern and western regions (Chan et al., 2017; Chan & Brown, 2019). We consider this proposed model to be incompatible with *Pteroptyx*, which occurs in mangroves and do not disperse across inland habitats to begin with. In this scenario, inland habitats represent ecological barriers and dispersal routes are consequently restricted to the coastline. However, the distribution of ancient mangroves are likely to be different compared to the present day and a recent study suggested that the dispersal of certain populations of *P. tener* in Peninsular Malaysia could have occurred along ancient river networks during the Pleistocene (Cheng, Munian, Sek-Aun, Faidi, & Ishak, 2019).

## CONCLUSIONS

Despite the charismatic attraction that fireflies hold in cultures all around the world, this study revealed that fundamental knowledge on Southeast Asian firefly biodiversity and evolutionary history is still lacking. More disconcertingly, much of their habitat, specifically forests and mangroves are suffering severe degradation and loss from pollution and land conversion (Jusoh & Hashim, 2012; Prasertkul, 2018; Wong & Yeap, 2012), setting the stage for an arms race to elucidate the true extent of firefly diversity before it is lost. Our data showed that *Pteroptyx* fireflies in Malaysia contain hitherto undiscovered cryptic and incipient species that are phylogeographically structured, thereby demonstrating the dynamic and complicated interplay between genes and the environment that drives, maintains, and distributes firefly biodiversity. Overall, this study contributes to the growing body of genetic work on fireflies and underscores the urgent need to increase the breadth and depth of geographic and genetic sampling to better understand the evolutionary history of fireflies.

## ACKNOWLEDGEMENTS

WFA thanks the Evolutionary Biology Lab at the National University of Singapore for support on molecular work.

## SUPPORTING INFORMATION

**Fig. S1.** Maximum likelihood phylogeny estimated using a partitioned analysis on a concatenated sequence matrix consisting of 2,119 bps of the *CO1* mitochondrial and CAD nuclear genes. Node values denote ultrafast bootstrap support.

**Fig. S2.** Bayesian phylogeney inferred from a concatenated sequence matrix consisting of 2,119 bps of the *CO1* mitochondrial and CAD nuclear genes. Node values denote posterior probabilities.

**Table S1.** List of genetic samples used in this study and their corresponding GenBank accession numbers.

**Supporting information 1.** Aligned sequences for the CO1 mitochondrial gene.

**Supporting information 2.** Aligned sequences for the CAD nuclear gene.

