## Supplementary figures and images for "Phylogeography and Molecular Species Delimitation Reveal Cryptic and Incipient Speciation in Synchronous Flashing Fireflies (Coleoptera: Lampyridae) of Southeast Asia"

### Figure S1

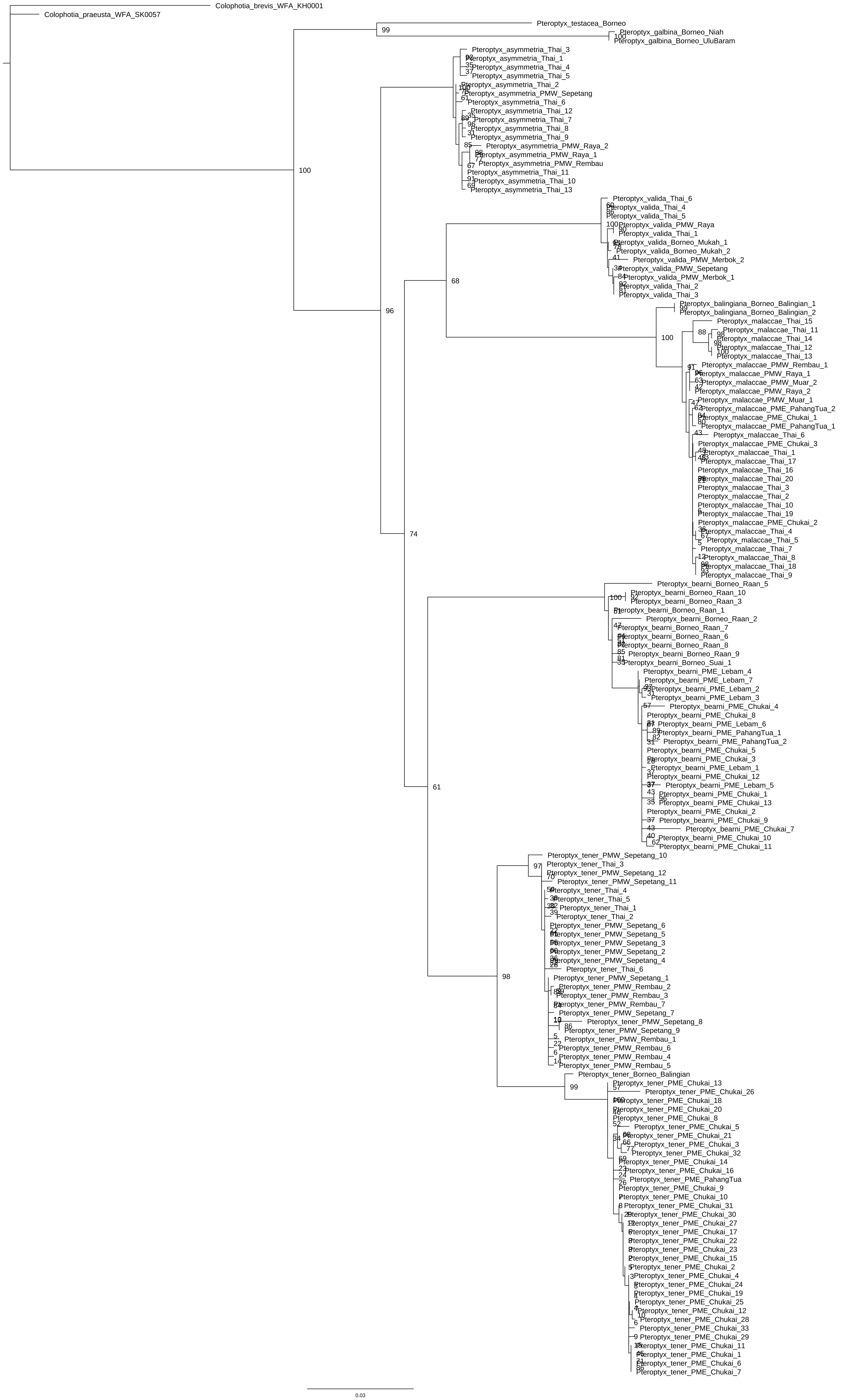

### Figure S2

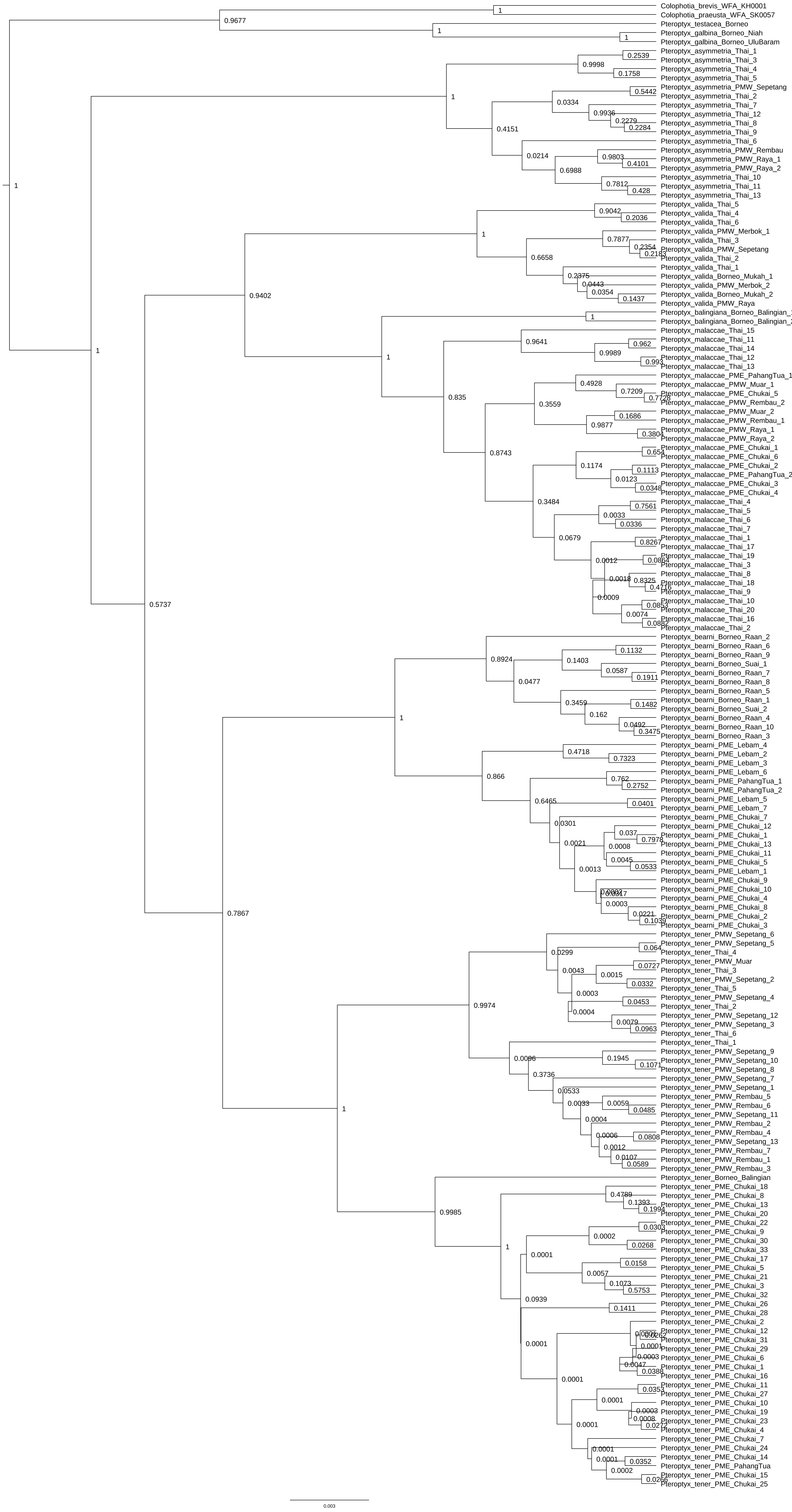
