## Supplementary material for "Phylogeography and Molecular Species Delimitation Reveal Cryptic and Incipient Speciation in Synchronous Flashing Fireflies (Coleoptera: Lampyridae) of Southeast Asia": Table S1

Table S1. List of DNA sequences used in this study

| **Sequence name** | **GenBank accession number according to molecular marker** | | | **Country** | **Locality** | **State** |
| --- | --- | --- | --- | --- | --- | --- |
|  | CAD (1-810) | CO1 fragment 1 (811-1475) | CO1 fragment 2 (1476-2119) |  |  |  |
| **OUTGROUP** |  |  |  |  |  |  |
| Colophotia_brevis_WFA_KH0001 | MF063271 | KY572911 | n.a | Malaysia | Merbok River | Kedah |
| Colophotia_praeusta_WFA_SK0057 | MF063272 | KY572909 | MF948237 | Malaysia | Rejang River | Sarawak |
| Pteroptyx_galbina_Borneo_UluBaram | n.a | KY572913 | n.a | Malaysia | Ulu Baram River | Sarawak |
| Pteroptyx_galbina_Borneo_Niah | MF063273 | KY572914 | MF948264 | Malaysia | Niah River | Sarawak |
| Pteroptyx_testacea_Borneo | MF063311 | KY572919 | MF948272 | Malaysia | Balingian River | Borneo |
| **INGROUP** |  |  |  |  |  |  |
| Pteroptyx_asymmetria_PMW_Raya_1 | MF063276 | KY572922 | MF948241 | Malaysia | Raya River | Negeri Sembilan |
| Pteroptyx_asymmetria_PMW_Raya_2 | n.a | KY572921 | n.a | Malaysia | Raya River | Negeri Sembilan |
| Pteroptyx_asymmetria_PMW_Rembau | MF063277 | KY572920 | MF948242 | Malaysia | Rembau River | Negeri Sembilan |
| Pteroptyx_asymmetria_PMW_Sepetang | MF063278 | KY572923 | MF948243 | Malaysia | Sepetang River | Perak |
| Pteroptyx_asymmetria_Thai_1 | n.a | MF624790 | n.a | Thailand | Laun, Ranong | Ranong |
| Pteroptyx_asymmetria_Thai_2 | n.a | MF624791 | n.a | Thailand | Laun, Ranong | Ranong |
| Pteroptyx_asymmetria_Thai_3 | n.a | MF624792 | n.a | Thailand | Laun, Ranong | Ranong |
| Pteroptyx_asymmetria_Thai_4 | n.a | MF624793 | n.a | Thailand | n.a. | n.a |
| Pteroptyx_asymmetria_Thai_5 | n.a | MF624794 | n.a | Thailand | n.a. | n.a |
| Pteroptyx_asymmetria_Thai_6 | n.a | MF624795 | n.a | Thailand | Takua Pa, PhangNga | PhangNga |
| Pteroptyx_asymmetria_Thai_7 | n.a | MF624796 | n.a | Thailand | n.a. | n.a |
| Pteroptyx_asymmetria_Thai_8 | n.a | MF624797 | n.a | Thailand | n.a. | n.a |
| Pteroptyx_asymmetria_Thai_9 | n.a | MF624798 | n.a | Thailand | Takua Pa, PhangNga | PhangNga |
| Pteroptyx_asymmetria_Thai_10 | n.a | MF624799 | n.a | Thailand | Takua Pa, PhangNga | PhangNga |
| Pteroptyx_asymmetria_Thai_11 | n.a | MF624800 | n.a | Thailand | PakBara, Satun | Satun |
| Pteroptyx_asymmetria_Thai_12 | n.a | MF624801 | n.a | Thailand | n.a. | n.a |
| Pteroptyx_asymmetria_Thai_13 | n.a | MF624802 | n.a | Thailand | Tapi River, Surat Thani | Surat Thani |
| Pteroptyx_valida_Borneo_Mukah_1 | n.a | KY573047 | n.a | Malaysia | Mukah River | Sarawak |
| Pteroptyx_valida_Borneo_Mukah_2 | MF063310 | KY573048 | MF948284 | Malaysia | Mukah River | Sarawak |
| Pteroptyx_valida_PMW_Merbok_1 | MF063308 | KY573050 | n.a | Malaysia | Merbok River | Kedah |
| Pteroptyx_valida_PMW_Merbok_2 | n.a | KY573049 | n.a | Malaysia | Merbok River | Kedah |
| Pteroptyx_valida_PMW_Raya | n.a | KY573046 | n.a | Malaysia | Raya River | Negeri Sembilan |
| Pteroptyx_valida_PMW_Sepetang | MF063309 | KY573051 | n.a | Malaysia | Sepetang River | Perak |
| Pteroptyx_valida_Thai_1 | n.a | MF624829 | n.a | Thailand | Muang, Nakhon Sri Thammarat | Nakhon Sri Thammarat |
| Pteroptyx_valida_Thai_2 | n.a | MF624830 | n.a | Thailand | Pak Bara, Satun | Satun |
| Pteroptyx_valida_Thai_3 | n.a | MF624831 | n.a | Thailand | n.a. | n.a |
| Pteroptyx_valida_Thai_4 | n.a | MF624832 | n.a | Thailand | Yeesan, Samut Songkhram | Samut Songkhram |
| Pteroptyx_valida_Thai_5 | n.a | MF624833 | n.a | Thailand | n.a. | n.a |
| Pteroptyx_valida_Thai_6 | n.a | MF624834 | n.a | Thailand | n.a. | n.a |
| Pteroptyx_balingiana_Borneo_Balingian_1 | MF063288 | KY572956 | MF948245 | Malaysia | Balingian River | Sarawak |
| Pteroptyx_balingiana_Borneo_Balingian_2 | MF063287 | KY572957 | MF948244 | Malaysia | Balingian River | Sarawak |
| Pteroptyx_bearni_Borneo_Raan_1 | MF063282 | KY572936 | MF948247 | Malaysia | Raan River | Sarawak |
| Pteroptyx_bearni_Borneo_Raan_2 | n.a | KY572927 | MF948246 | Malaysia | Raan River | Sarawak |
| Pteroptyx_bearni_Borneo_Raan_3 | n.a | KY572939 | MF948249 | Malaysia | Raan River | Sarawak |
| Pteroptyx_bearni_Borneo_Raan_4 | MF063285 | n.a | n.a | Malaysia | Raan River | Sarawak |
| Pteroptyx_bearni_Borneo_Raan_5 | n.a | KY572932 | n.a | Malaysia | Raan River | Sarawak |
| Pteroptyx_bearni_Borneo_Raan_6 | n.a | KY572926 | n.a | Malaysia | Raan River | Sarawak |
| Pteroptyx_bearni_Borneo_Raan_7 | n.a | KY572948 | n.a | Malaysia | Raan River | Sarawak |
| Pteroptyx_bearni_Borneo_Raan_8 | n.a | KY572950 | n.a | Malaysia | Raan River | Sarawak |
| Pteroptyx_bearni_Borneo_Raan_9 | n.a | KY572952 | n.a | Malaysia | Raan River | Sarawak |
| Pteroptyx_bearni_Borneo_Raan_10 | n.a | n.a | MF948248 | Malaysia | Raan River | Sarawak |
| Pteroptyx_bearni_Borneo_Suai_1 | MF063283 | KY572937 | n.a | Malaysia | Suai River | Sarawak |
| Pteroptyx_bearni_Borneo_Suai_2 | MF063284 | n.a | n.a | Malaysia | Suai River | Sarawak |
| Pteroptyx_bearni_PME_Chukai_1 | MF063286 | KY572934 | MF948259 | Malaysia | Chukai River | Terengganu |
| Pteroptyx_bearni_PME_Chukai_2 | MF063298 | KY572946 | MF948252 | Malaysia | Chukai River | Terengganu |
| Pteroptyx_bearni_PME_Chukai_3 | n.a | KY572925 | MF948251 | Malaysia | Chukai River | Terengganu |
| Pteroptyx_bearni_PME_Chukai_4 | n.a | KY572929 | MF948253 | Malaysia | Chukai River | Terengganu |
| Pteroptyx_bearni_PME_Chukai_5 | n.a | KY572930 | MF948257 | Malaysia | Chukai River | Terengganu |
| Pteroptyx_bearni_PME_Chukai_7 | n.a | KY572933 | MF948258 | Malaysia | Chukai River | Terengganu |
| Pteroptyx_bearni_PME_Chukai_8 | n.a | KY572947 | n.a | Malaysia | Chukai River | Terengganu |
| Pteroptyx_bearni_PME_Chukai_9 | n.a | KY572940 | MF948255 | Malaysia | Chukai River | Terengganu |
| Pteroptyx_bearni_PME_Chukai_10 | n.a | KY572941 | MF948256 | Malaysia | Chukai River | Terengganu |
| Pteroptyx_bearni_PME_Chukai_11 | n.a | KY572942 | n.a | Malaysia | Chukai River | Terengganu |
| Pteroptyx_bearni_PME_Chukai_12 | n.a | KY572945 | n.a | Malaysia | Chukai River | Terengganu |
| Pteroptyx_bearni_PME_Chukai_13 | n.a | n.a | MF948254 | Malaysia | Chukai River | Terengganu |
| Pteroptyx_bearni_PME_Lebam_1 | MF063279 | KY572938 | MF948262 | Malaysia | Lebam River | Johor |
| Pteroptyx_bearni_PME_Lebam_2 | n.a | KY572935 | MF948260 | Malaysia | Lebam River | Johor |
| Pteroptyx_bearni_PME_Lebam_3 | n.a | KY572931 | MF948261 | Malaysia | Lebam River | Johor |
| Pteroptyx_bearni_PME_Lebam_4 | n.a | KY572944 | n.a | Malaysia | Lebam River | Johor |
| Pteroptyx_bearni_PME_Lebam_5 | MF063280 | KY572949 | n.a | Malaysia | Lebam River | Johor |
| Pteroptyx_bearni_PME_Lebam_6 | n.a | KY572953 | n.a | Malaysia | Lebam River | Johor |
| Pteroptyx_bearni_PME_Lebam_7 | n.a | n.a | MF948263 | Malaysia | Lebam River | Johor |
| Pteroptyx_bearni_PME_PahangTua_1 | MF063281 | KY572924 | n.a | Malaysia | Pahang Tua River | Pahang |
| Pteroptyx_bearni_PME_PahangTua_2 | n.a | KY572954 | n.a | Malaysia | Pahang Tua River | Pahang |
| Pteroptyx_malaccae_PME_Chukai_1 | MF063293 | KY572963 | n.a | Malaysia | Chukai River | Terengganu |
| Pteroptyx_malaccae_PME_Chukai_2 | MF063297 | n.a | MF948268 | Malaysia | Chukai River | Terengganu |
| Pteroptyx_malaccae_PME_Chukai_3 | n.a | n.a | MF948271 | Malaysia | Chukai River | Terengganu |
| Pteroptyx_malaccae_PME_Chukai_4 | MF063294 | n.a | n.a | Malaysia | Chukai River | Terengganu |
| Pteroptyx_malaccae_PME_Chukai_5 | MF063295 | n.a | n.a | Malaysia | Chukai River | Terengganu |
| Pteroptyx_malaccae_PME_Chukai_6 | MF063296 | n.a | n.a | Malaysia | Chukai River | Terengganu |
| Pteroptyx_malaccae_PME_PahangTua_1 | MF063289 | KY572965 | MF948265 | Malaysia | Pahang Tua River | Pahang |
| Pteroptyx_malaccae_PME_PahangTua_2 | n.a | KY572964 | MF948267 | Malaysia | Pahang Tua River | Pahang |
| Pteroptyx_malaccae_PMW_Muar_1 | MF063290 | KY572962 | MF948269 | Malaysia | Muar River | Johor |
| Pteroptyx_malaccae_PMW_Muar_2 | n.a | KY572958 | MF948266 | Malaysia | Muar River | Johor |
| Pteroptyx_malaccae_PMW_Raya_1 | n.a | KY572959 | n.a | Malaysia | Raya River | Negeri Sembilan |
| Pteroptyx_malaccae_PMW_Raya_2 | n.a | KY572960 | n.a | Malaysia | Raya River | Negeri Sembilan |
| Pteroptyx_malaccae_PMW_Rembau_1 | MF063292 | KY572961 | MF948270 | Malaysia | Rembau River | Negeri Sembilan |
| Pteroptyx_malaccae_PMW_Rembau_2 | MF063291 | n.a | n.a | Malaysia | Rembau River | Negeri Sembilan |
| Pteroptyx_malaccae_Thai_1 | n.a | MF624803 | n.a | Thailand | Tapi river branch, Surat Thani | |
| Pteroptyx_malaccae_Thai_2 | n.a | MF624804 | n.a | Thailand | Bang Saphan, Prachuap Khiri Khan | |
| Pteroptyx_malaccae_Thai_3 | n.a | MF624805 | n.a | Thailand | n.a. | n.a |
| Pteroptyx_malaccae_Thai_4 | n.a | MF624806 | n.a | Thailand | n.a. | n.a |
| Pteroptyx_malaccae_Thai_5 | n.a | MF624807 | n.a | Thailand | n.a. | n.a |
| Pteroptyx_malaccae_Thai_6 | n.a | MF624808 | n.a | Thailand | Yeesan, Samut Songkhram | Samut Songkhram |
| Pteroptyx_malaccae_Thai_7 | n.a | MF624809 | n.a | Thailand | n.a. | n.a |
| Pteroptyx_malaccae_Thai_8 | n.a | MF624810 | n.a | Thailand | n.a. | n.a |
| Pteroptyx_malaccae_Thai_9 | n.a | MF624811 | n.a | Thailand | n.a. | n.a |
| Pteroptyx_malaccae_Thai_10 | n.a | MF624812 | n.a | Thailand | n.a. | n.a |
| Pteroptyx_malaccae_Thai_11 | n.a | MF624813 | n.a | Thailand | Ban Nam Chiao, Trat | Trat |
| Pteroptyx_malaccae_Thai_12 | n.a | MF624814 | n.a | Thailand | Ban Nam Chiao, Trat | Trat |
| Pteroptyx_malaccae_Thai_13 | n.a | MF624815 | n.a | Thailand | n.a. | n.a |
| Pteroptyx_malaccae_Thai_14 | n.a | MF624816 | n.a | Thailand | n.a. | n.a |
| Pteroptyx_malaccae_Thai_15 | n.a | MF624817 | n.a | Thailand | Weru Wet land, Chanthaburi | Chanthaburi |
| Pteroptyx_malaccae_Thai_16 | n.a | MF624818 | n.a | Thailand | n.a. |  |
| Pteroptyx_malaccae_Thai_17 | n.a | MF624819 | n.a | Thailand | Tapi river branch, Surat Thani | Surat Thani |
| Pteroptyx_malaccae_Thai_18 | n.a | MF624820 | n.a | Thailand | n.a. | n.a |
| Pteroptyx_malaccae_Thai_19 | n.a | MF624821 | n.a | Thailand | n.a. | n.a |
| Pteroptyx_malaccae_Thai_20 | n.a | MF624822 | n.a | Thailand | n.a. | n.a |
| Pteroptyx_tener_Borneo_Balingian | n.a | KY572989 | MF948282 | Malaysia | Balingian River | Sarawak |
| Pteroptyx_tener_PME_Chukai_1 | MF063305 | KY572985 | MF948276 | Malaysia | Chukai River | Terengganu |
| Pteroptyx_tener_PME_Chukai_2 | MF063307 | KY572991 | MF948274 | Malaysia | Chukai River | Terengganu |
| Pteroptyx_tener_PME_Chukai_3 | n.a | KY572975 | MF948273 | Malaysia | Chukai River | Terengganu |
| Pteroptyx_tener_PME_Chukai_4 | MF063306 | KY573001 | n.a | Malaysia | Chukai River | Terengganu |
| Pteroptyx_tener_PME_Chukai_5 | n.a | KY573016 | MF948275 | Malaysia | Chukai River | Terengganu |
| Pteroptyx_tener_PME_Chukai_6 | MF063299 | n.a | MF948277 | Malaysia | Chukai River | Terengganu |
| Pteroptyx_tener_PME_Chukai_7 | MF063300 | n.a | MF948283 | Malaysia | Chukai River | Terengganu |
| Pteroptyx_tener_PME_Chukai_8 | n.a | KY572966 | n.a | Malaysia | Chukai River | Terengganu |
| Pteroptyx_tener_PME_Chukai_9 | n.a | KY572967 | n.a | Malaysia | Chukai River | Terengganu |
| Pteroptyx_tener_PME_Chukai_10 | n.a | KY572968 | n.a | Malaysia | Chukai River | Terengganu |
| Pteroptyx_tener_PME_Chukai_11 | n.a | KY572983 | n.a | Malaysia | Chukai River | Terengganu |
| Pteroptyx_tener_PME_Chukai_12 | n.a | KY572984 | n.a | Malaysia | Chukai River | Terengganu |
| Pteroptyx_tener_PME_Chukai_13 | n.a | KY572987 | n.a | Malaysia | Chukai River | Terengganu |
| Pteroptyx_tener_PME_Chukai_14 | n.a | KY572990 | n.a | Malaysia | Chukai River | Terengganu |
| Pteroptyx_tener_PME_Chukai_15 | n.a | KY572992 | n.a | Malaysia | Chukai River | Terengganu |
| Pteroptyx_tener_PME_Chukai_16 | n.a | KY572994 | n.a | Malaysia | Chukai River | Terengganu |
| Pteroptyx_tener_PME_Chukai_17 | n.a | KY572995 | n.a | Malaysia | Chukai River | Terengganu |
| Pteroptyx_tener_PME_Chukai_18 | n.a | KY572997 | n.a | Malaysia | Chukai River | Terengganu |
| Pteroptyx_tener_PME_Chukai_19 | n.a | KY572999 | n.a | Malaysia | Chukai River | Terengganu |
| Pteroptyx_tener_PME_Chukai_20 | n.a | KY573000 | n.a | Malaysia | Chukai River | Terengganu |
| Pteroptyx_tener_PME_Chukai_21 | n.a | KY573002 | n.a | Malaysia | Chukai River | Terengganu |
| Pteroptyx_tener_PME_Chukai_22 | n.a | KY573004 | n.a | Malaysia | Chukai River | Terengganu |
| Pteroptyx_tener_PME_Chukai_23 | n.a | KY573005 | n.a | Malaysia | Chukai River | Terengganu |
| Pteroptyx_tener_PME_Chukai_24 | n.a | KY573008 | n.a | Malaysia | Chukai River | Terengganu |
| Pteroptyx_tener_PME_Chukai_25 | n.a | KY573013 | n.a | Malaysia | Chukai River | Terengganu |
| Pteroptyx_tener_PME_Chukai_26 | n.a | KY573015 | n.a | Malaysia | Chukai River | Terengganu |
| Pteroptyx_tener_PME_Chukai_27 | n.a | KY573017 | n.a | Malaysia | Chukai River | Terengganu |
| Pteroptyx_tener_PME_Chukai_28 | n.a | KY573018 | n.a | Malaysia | Chukai River | Terengganu |
| Pteroptyx_tener_PME_Chukai_29 | n.a | KY573020 | n.a | Malaysia | Chukai River | Terengganu |
| Pteroptyx_tener_PME_Chukai_30 | n.a | KY573021 | n.a | Malaysia | Chukai River | Terengganu |
| Pteroptyx_tener_PME_Chukai_31 | n.a | KY573030 | n.a | Malaysia | Chukai River | Terengganu |
| Pteroptyx_tener_PME_Chukai_32 | n.a | KY573037 | n.a | Malaysia | Chukai River | Terengganu |
| Pteroptyx_tener_PME_Chukai_33 | n.a | KY573040 | n.a | Malaysia | Chukai River | Terengganu |
| Pteroptyx_tener_PME_PahangTua | n.a | KY573044 | n.a | Malaysia | Pahang Tua River | Pahang |
| Pteroptyx_tener_PMW_Muar | MF063301 | n.a | n.a | Malaysia | Muar River | Johor |
| Pteroptyx_tener_PMW_Rembau_1 | n.a | KY573026 | n.a | Malaysia | Rembau River | Negeri Sembilan |
| Pteroptyx_tener_PMW_Rembau_2 | n.a | KY573027 | n.a | Malaysia | Rembau River | Negeri Sembilan |
| Pteroptyx_tener_PMW_Rembau_3 | n.a | KY572986 | n.a | Malaysia | Rembau River | Negeri Sembilan |
| Pteroptyx_tener_PMW_Rembau_4 | n.a | KY573007 | n.a | Malaysia | Rembau River | Negeri Sembilan |
| Pteroptyx_tener_PMW_Rembau_5 | n.a | KY573010 | n.a | Malaysia | Rembau River | Negeri Sembilan |
| Pteroptyx_tener_PMW_Rembau_6 | n.a | KY572973 | n.a | Malaysia | Rembau River | Negeri Sembilan |
| Pteroptyx_tener_PMW_Rembau_7 | n.a | KY572976 | n.a | Malaysia | Rembau River | Negeri Sembilan |
| Pteroptyx_tener_PMW_Sepetang_1 | MF063304 | KY572970 | MF948281 | Malaysia | Sepetang River | Perak |
| Pteroptyx_tener_PMW_Sepetang_2 | n.a | KY572969 | n.a | Malaysia | Sepetang River | Perak |
| Pteroptyx_tener_PMW_Sepetang_3 | n.a | KY572971 | n.a | Malaysia | Sepetang River | Perak |
| Pteroptyx_tener_PMW_Sepetang_4 | n.a | KY572972 | n.a | Malaysia | Sepetang River | Perak |
| Pteroptyx_tener_PMW_Sepetang_5 | n.a | KY572977 | n.a | Malaysia | Sepetang River | Perak |
| Pteroptyx_tener_PMW_Sepetang_6 | n.a | KY572978 | n.a | Malaysia | Sepetang River | Perak |
| Pteroptyx_tener_PMW_Sepetang_7 | n.a | KY572979 | n.a | Malaysia | Sepetang River | Perak |
| Pteroptyx_tener_PMW_Sepetang_8 | n.a | KY572993 | n.a | Malaysia | Sepetang River | Perak |
| Pteroptyx_tener_PMW_Sepetang_9 | n.a | KY573011 | n.a | Malaysia | Sepetang River | Perak |
| Pteroptyx_tener_PMW_Sepetang_10 | n.a | n.a | MF948278 | Malaysia | Sepetang River | Perak |
| Pteroptyx_tener_PMW_Sepetang_11 | n.a | n.a | MF948279 | Malaysia | Sepetang River | Perak |
| Pteroptyx_tener_PMW_Sepetang_12 | MF063303 | n.a | MF948280 | Malaysia | Sepetang River | Perak |
| Pteroptyx_tener_PMW_Sepetang_13 | MF063302 | n.a | n.a | Malaysia | Sepetang River | Perak |
| Pteroptyx_tener_Thai_1 | n.a | MF624823 | n.a | Thailand | Tapi River branch | Surat Thani |
| Pteroptyx_tener_Thai_2 | n.a | MF624824 | n.a | Thailand | Tapi River branch | Surat Thani |
| Pteroptyx_tener_Thai_3 | n.a | MF624825 | n.a | Thailand | Tapi River branch | Surat Thani |
| Pteroptyx_tener_Thai_4 | n.a | MF624826 | n.a | Thailand | Tapi River branch | Surat Thani |
| Pteroptyx_tener_Thai_5 | n.a | MF624827 | n.a | Thailand | Tapi River branch | Surat Thani |
| Pteroptyx_tener_Thai_6 | n.a | MF624828 | n.a | Thailand | n.a. | n.a |
